# Symbiotic acacia ants drive nesting behavior by birds in an African savanna

**DOI:** 10.1101/2023.03.06.531340

**Authors:** Lujan Ema, Ryen Nielsen, Zoe Short, Samuel Wicks, Wilson Nderitu Watetu, Leo M. Khasoha, Todd M. Palmer, Jacob R. Goheen, Jesse M. Alston

**Author notes:** These authors contributed equally to this work (Lujan Ema, Ryen Nielsen).

## Abstract

Mutualisms between plants and ants are common features of tropical ecosystems around the globe and can have cascading effects on interactions with the ecological communities in which they occur. In an African savanna, we assessed whether acacia ants influence nest site selection by tree-nesting birds. Birds selected nest sites in trees inhabited by ant species that vigorously defend against browsing mammals. Future research could address the extent to which hatching and fledging rates depend on the species of ant symbiont, and why ants tolerate nesting birds but not other tree associates (especially insects).

---

Mutualisms structure biodiversity and ecosystem function (Stachowicz 2001). Mutualisms between plants and ants are particularly widespread across the tropics (Christian 2001, Frederickson *et al*. 2005, Palmer & Brody 2013, Prior *et al*. 2015), including the savannas of East Africa (Young *et al*. 1996, Stanton & Palmer 2011, Palmer & Brody 2013, Hays *et al*.2022). In such ecosystems, whistling thorn trees (*Acacia [Vachellia] drepanolobium*) form a near-monoculture, comprising 95-99% of the canopy layer (Young *et al*. 1996). Four ant species (*Crematogaster mimosae, C. nigriceps, C. sjostedti,* and *Tetraponera penzigi*) are symbionts of whistling thorn trees, which produce extrafloral nectar and swollen-thorn domatia to recruit and maintain colonies (Palmer *et al*. 2008).

Ant species exclusively occupy host trees, with a single species typically controlling the canopies of individual trees at any given time. Further, the four ant species vary in the benefits they provide and costs they impose to host trees. *Crematogaster mimosae* and *C. nigriceps* aggressively defend trees against mammalian and insect herbivores, and they are particularly effective at deterring catastrophic (lethal) herbivory by elephants (*Loxodonta africana*; Goheen & Palmer 2010, Palmer & Brody 2013). By sterilizing its host trees, *C. nigriceps* additionally functions as a short-term (one to several years) parasite, but it enhances lifetime fitness by offering protection to otherwise vulnerable, pre-reproductive trees (Stanton *et al*. 1999, Palmer *et al*. 2010). In contrast, *T. penzigi* and *C. sjostedti* provide only moderate to minimal protection, respectively, against herbivory (Palmer & Brody 2007, Palmer *et al*. 2010).

Despite the ants’ presence, several bird species—such as gray-capped social weavers (*Pseudonigrita arnaudi*), gray-headed sparrows (*Passer griseus*), and superb starlings (*Lamprotornis superbus*)—often nest in whistling thorn trees. Although birds nest in ant-defended acacias in Central America (Janzen 1969, Young *et al*. 1990, Flaspohler & Laska 1994, Oliveras de Ita & Rojas-Soto 2006), ants are nest predators (Smith *et al*. 2007, Menezes & Marini 2017) and can deter birds from feeding in ant-defended trees (Haemig 1994, Aho *et al*.1997, Philpott *et al*. 2005). These contrasting observations from the Neotropics generate distinct predictions regarding whether and how birds distinguish among host trees occupied by different ant symbionts. If acacia ants defend against all disturbances to host trees, then birds should select trees occupied by less aggressive symbionts (i.e., *C. sjostedti* and *T. penzigi*, to a lesser extent) for nesting. However, it also is possible that acacia ants confer protection to bird nests, in which case birds should select for trees with aggressive symbionts (i.e., *C. mimosae* and *C. nigriceps)*.

To uncover associations between birds and acacia ants, we systematically searched for bird nests in whistling thorn savannas at Mpala Research Centre and Conservancy (0°17’ N, 36°53’ E), Laikipia, Kenya in June 2022. We identified species by nest architecture: gray-capped social weavers build spherical nests with bottom-facing entrances (usually with multiple nests in the same tree), while superb starlings and gray-headed sparrows build gourd-shaped nests with side-facing entrances (usually with one nest per tree). Nests of superb starlings and gray-headed sparrows can be distinguished by the size of the entrance (starling nests have entrances large enough to fit a hand into, while entrances of sparrow nests are smaller). For each “used” tree in which we found bird nests, we identified the four nearest neighbors above 0.5 m tall, classifying these as “available”. For both used and available trees, we measured tree height, diameter at 30 cm from the base, whether the tree was alive or dead, canopy area (calculated by measuring the width and length of the canopy and estimating its area, where area =π * width * length), and the species of ant symbiont occupying the tree. We performed logistic regression to quantify the influence of these predictors on nest tree selection and calculated variance inflation factors to check for multicollinearity among predictors.

Some of these ant species (particularly C. *sjostedti* and *C. mimosae*) are spatially clustered on the landscape and inordinately likely to inhabit neighboring trees. To ensure that such spatial autocorrelation did not bias our results, we also conducted a second logistic regression in which we substituted the available trees we described above for a new set of available trees that were < 0.5 m tall and located within 10 m of glades (nutrient-rich, open grazing lawns that form after livestock graze the area for an extended period of time) or termite mounds. Nests were typically found within or near these landscape features. This second set of available trees was surveyed for previous research (Palmer *et al*. 2010), and data were only available for height and ant species occupant.

We used the ‘car’ package (v3.0.12; Fox & Weisberg 2019) to calculate variance inflation factors and the R statistical software environment (v4.2.1; R Core Team 2020) to perform all statistical analyses.

We located 60 nests in total (34 superb starling, 16 gray-headed sparrow, 8 gray-capped social weaver, and 2 cup nests created by unknown species). Of these nests, 45 were located in trees inhabited by *Crematogaster mimosae*, 14 in trees inhabited by *Crematogaster nigriceps*, and 1 in a tree inhabited by *Crematogaster sjostedti*. All nests were in live trees that were more than 1.5 m in height.

Our first logistic regression model identified height (*β* = 0.002; *p* < 0.0001) and occupancy by *C. nigriceps* (*β* = 0.17; *p* < 0.01) as the most important predictors of nest selection (Table 1; Fig. 1A). For each 1 m increase in height, the odds that birds nested in a tree increased by 20% (95% CI: 12 – 29%). The odds that birds nested in a tree inhabited by *C. nigriceps* were 18% higher than for those inhabited by *C. mimosae* (the reference category; 95% CI: 5 – 34%). Other predictors were not significant.

**Table 1.** Coefficient estimates in the logistic regression model and 95% confidence intervals. Bold variables denote significance at α = 0.05.

| **Model 1** |  |  |  | **Model 2** |  |  |  |
| --- | --- | --- | --- | --- | --- | --- | --- |
| *Variable* | *Estimate* | *LCL* | *UCL* | *Variable* | *Estimate* | *LCL* | *UCL* |
| ***Intercept*** | **-0.134** | **-0.254** | **-0.014** | ***Intercept*** | **0.231** | **0.098** | **0.364** |
| *Live* | -0.064 | -0.399 | 0.270 | *Height* | 0.038 | -0.006 | 0.083 |
| ***Height*** | **0.183** | **0.115** | **0.251** | *C. nigriceps* | 0.120 | -0.031 | 0.271 |
| *Canopy Area* | -0.014 | -0.029 | 0.002 | ***C. sjostedti*** | **-0.311** | **-0.429** | **-0.192** |
| ***C. nigriceps*** | **0.169** | **0.046** | **0.292** | ***T. penzigi*** | **-0.299** | **-0.465** | **-0.134** |
| *C. sjostedti* | -0.168 | -0.357 | 0.021 | *C. sjostedti + T. penzigi* | -0.319 | -1.086 | 0.449 |
| *T. penzigi* | -0.101 | -0.352 | 0.150 | *No ants* | -0.256 | -0.654 | 0.142 |
| *No ants* | -0.082 | -0.386 | 0.222 |  |  |  |  |

**Figure 1.**
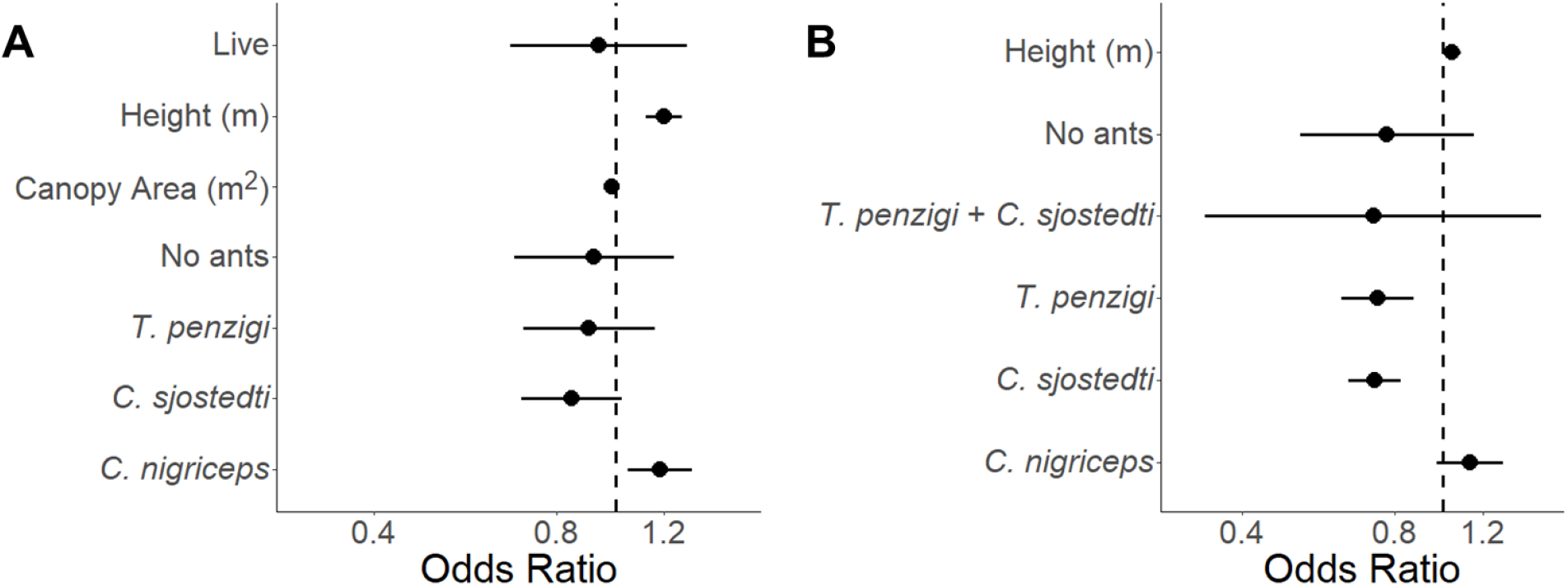
Unscaled odds ratios associated with each variable in the nest site selection model. Error bars represent 95% confidence intervals. The results of Model 1 are shown in Panel A; the results of Model 2 are shown in Panel B.

Our second logistic regression model did not identify height as an important predictor of nest selection (*β* = 0.04; *p* = 0.09), but it did identify ant species as an important predictor. Compared to trees inhabited by *C. mimosae*, the odds that trees inhabited by *C. nigriceps* contained nests were roughly equal (95% CI: −3 – 34%), but the odds that trees inhabited by *C. sjostedti* contained nests were 27% lower (95% CI: 17 – 35%), and the odds that trees inhabited by *T. penzigi* contained nests were 26% lower (95% CI: 13 – 37%).

Birds almost always nested in trees inhabited by aggressive defenders of host trees (*C. nigriceps* and *C. mimosae*), particularly *C. nigriceps* (Table 1, Fig. 1). The selection of nesting sites inhabited by more aggressive ant species may reduce risk of nest predation (Young *et al*.1990), which can reduce lifetime fitness in birds (Freeman et al. 2020; Martin 1993). Future research with longitudinal data on nest survival may elucidate the fitness benefits of ant symbionts for birds.

Tree architecture likely plays an important role in nest site selection by birds. In addition to protecting whistling thorn trees from herbivory, some acacia ants change the architecture of *A. drepanolobium*. Because *Crematogaster nigriceps* is an inferior competitor to other *Crematogaster* spp., it prunes apical buds, which shortens shoots and reduces the likelihood of contact with host trees occupied by *C. nigriceps* and *C. sjostedti* (Stanton *et al*. 1999). As such, occupancy by *C. nigriceps* results in substantially denser canopies, which likely provide concealment and further protection from predators (see also Oliveras de Ita & Rojas-Soto 2006, Latif *et al*. 2012), which include snakes, mesocarnivores, and raptors (W. Watetu, pers. obs.).

Although *C. mimosae* and *C. nigriceps* vigorously defend their host trees from both vertebrate and invertebrate herbivores, and some ants are nest predators (e.g., Suarez *et al*. 2005, Smith *et al*. 2007, Menezes & Marini 2017), *C. mimosae* and *C. nigriceps* apparently attack neither nestlings nor adult birds (W. Watetu, pers. obs.). Acacia ants can distinguish between wind-induced and herbivore-induced vibrations (Hager & Krausa 2019), but ants readily attack humans manipulating bird nests. The cues ants use to differentiate birds from herbivores against which they defend trees remain unclear, but our study suggests that either ants can distinguish between sources of vibrations even better than is currently appreciated, that other cues (e.g., chemical or visual cues) may also trigger ant defense of trees, or that bird nests have chemical or structural characteristics that deter ants from entering them.

Birds are not the only occupants of *A. drepanolobium*, and acacia ants may influence the ecology of other *A. drepanolobium* inhabitants as well. Several arboreal reptiles inhabit *A. drepanolobium*, and the most common of these (*Lygodactylus keniensis*, a gecko) selects for trees inhabited by the least aggressive ant, *C. sjostedti* (Pringle *et al*. 2015), perhaps because the more aggressive ant species inhibit the elephant damage that creates the gecko’s preferred microhabitats (Pringle 2008). It is possible that acacia ants similarly influence habitat selection by the other, less common arboreal reptiles in this system, by directly defending trees from animals moving in them, by influencing patterns of tree damage and herbivory by large herbivores, or by altering the architecture of tree canopies. Ants could likewise influence the use of *A. drepanolobium* by the other animals known to inhabit these trees, including other bird species, primates, and invertebrates.

In summary, birds in an East African whistling thorn savanna select nest sites in trees defended by the most aggressive acacia ants, particularly a species (*C. nigriceps*) that alters tree architecture such that the canopy is denser. This raises questions for future work: Are birds selecting nest trees based on the aggressiveness of ant symbionts *per se*, correlates of ant symbionts (like the denser architecture of trees inhabited by *C. nigriceps*), or both? Do ant symbionts differentially affect hatching and fledging rates? How do ants distinguish between birds and other animals they defend trees against? Do the acacia ants benefit other animal species that also inhabit these trees? Further research to answer these questions may reveal much more about how mutualisms operate, and cascading effects for other species in interaction webs.

## Acknowledgments

We thank Mpala Research Centre for hosting the field course during which we collected data and A. Klass for assisting with field data collection. Financial support was provided by the University of Wyoming’s Center for Global Studies, the University of Wyoming’s Biodiversity Institute and a National Science Foundation grant to TMP (DEB 1149980).

## Data Availability Statement

All data and code underlying the analyses detailed in this manuscript will be archived in *Zenodo* upon acceptance.

## Author Contributions

JRG conceived the ideas; EL, RN, and JA led the drafting of the manuscript; EL, RN, ZS, SW, and TMP collected field data; JA led statistical analyses. All authors contributed critically to drafts of the manuscript and gave final approval for publication.

